# Mechanical frustration of phase separation in the cell nucleus by chromatin

**DOI:** 10.1101/2020.12.24.424222

**Authors:** Yaojun Zhang, Daniel S.W. Lee, Yigal Meir, Clifford P. Brangwynne, Ned S. Wingreen

**Author notes:** These two authors contributed equally (Yaojun Zhang, Daniel S.W. Lee).

## Abstract

Liquid-liquid phase separation is a fundamental mechanism underlying subcellular organization. Motivated by the striking observation that optogenetically-generated droplets in the nucleus display suppressed coarsening dynamics, we study the impact of chromatin mechanics on droplet phase separation. We combine theory and simulation to show that crosslinked chromatin can mechanically suppress droplets’ coalescence and ripening, as well as quantitatively control their number, size, and placement. Our results highlight the role of the subcellular mechanical environment on condensate regulation.

---

Eukaryotic cells are host to a multiplicity of membraneless compartments, many of which form and dissolve as needed to enable central cellular functions – from ribosome assembly to transcription, signaling, and metabolism [1,2]. These compartments form via liquid-liquid phase separation, driven by multivalent interactions among proteins and/or RNAs [3–6]. Unlike conventional phase separation, e.g. the demixing of oil and water, biomolecular phase separation takes place in the complex environment of the cell: the cytoplasm is scaffolded by a dynamic cytoskeletal network, while the nucleus is packed with viscoelastic chromatin [7,8]. How do such complex environments impact the equilibrium states and out-of-equilibrium dynamics of biomolecular condensates?

The thermodynamic ground state of two immiscible liquids is a single droplet of one liquid immersed in the other. Natural and synthetic condensates in cells, however, typically appear as dispersed droplets [9–11]. To test the stability of droplets in the cell, we used a novel optogenetic system [12,13] to create droplets in the nucleus (Fig. 1). We used patterned local activation to create ∼ 10 large droplets, followed by global activation to generate many small droplets. The areas of the initially created large droplets (Fig. 1(b)) are shown as functions of time (Fig. 1(c)). See Supplemental Material for details [14].

**FIG. 1.**
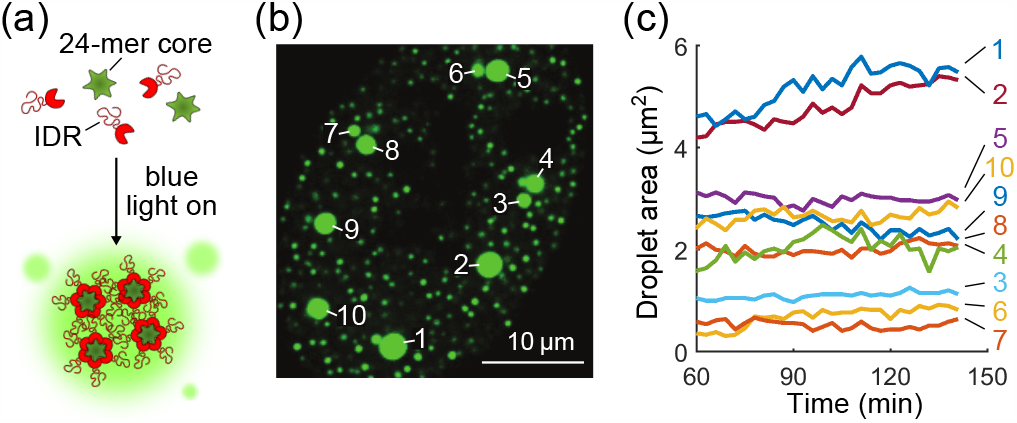
Droplet coarsening is suppressed in the cell nucleus. (a) Schematic of light-activated intracellular phase separation. Upon blue-light illumination, up to 24 intrinsically disordered protein regions (IDRs) bind to each 24-mer core. These subsequently phase separate due to multivalent IDR interactions. (b) Optogenetically-generated fluorescent protein droplets (green) in the nucleus of a U2OS cell. Large droplets (labeled with numbers) are created by patterned local activation. Subsequently, many much smaller droplets are created by global activation. (The large droplets generated this way initially have higher IDR-to-core ratios than the small droplets due to a diffusive capture mechanism [13], and droplets change their sizes as this ratio equilibrates; we therefore focus on the time evolution of droplet sizes after this effect subsides.) (c) Time evolution of areas of large droplets in (b), starting at 60 min after global activation. (Full time-course trajectories are shown in Fig. S1.)

Given the drastic size difference between the large and small droplets, naively, one would expect small droplets to lose material to large ones, in a process known as Ostwald ripening [15]. Theoretically, for a large droplet of radius *R*_1_ surrounded by many small droplets each of radius *R*_2_ all at a distance *L*, the cube of radius of the large droplet is predicted to grow as [14]:

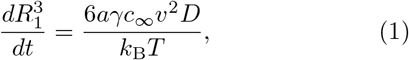

where *a* = (*R*_1_*/R*_2_ − 1)*/*(1 − *R*_1_*/L*) is a geometrical factor, *γ* is the surface tension, *c*_*∞*_ is the solubility of the droplet molecules, *v* is their molecular volume, *D* is their diffusion coefficient, *k*_B_ is the Boltzmann constant, and *T* is temperature (we set *T* = 300 K here and in our simulations). Taking *R*_1_ = 1 *µ*m, *R*_2_ = 0.2 *µ*m, *L* = 3 *µ*m, and using the estimated biological parameters in Table I [10,13], Eq. (1) predicts 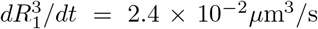. Therefore, within just ∼1 minute we would expect large droplets to double in volume. However, Fig. 1(c) shows that droplets 1 and 2 have mild growth, and the rest of the large droplets barely change in size over 80 minutes. Based on the growth rate of droplets 1 and 2 we estimate the upper bound of the ripening rate to be 2×10^*−*4^*µ*m^3^*/*s. It thus seems that the coarsening dynamics of droplets is strongly suppressed.

**TABLE I.** Ostwald ripening parameters and rates for experiment and for simulations of phase separation without chromatin, Case (i), and with uncrosslinked chromatin, Case (ii).

| | $\gamma$ | $c_\infty$ | $v$ | $D$ | rate<sup>(t)</sup> | rate<sup>(m)</sup> |
| --- | --- | --- | --- | --- | --- | --- |
| | N/m | 1/$\mu\text{m}^3$ | $\text{nm}^3$ | $\mu\text{m}^2/\text{s}$ | $\mu\text{m}^3/\text{s}$ | $\mu\text{m}^3/\text{s}$ |
| Exp | $5 \times 10^{-7}$ | 1200 | $6.7 \times 10^4$ | 1 | $2.4 \times 10^{-2}$ | $\leq 2 \times 10^{-4}$ |
| (i) | $3.8 \times 10^{-6}$ | 271 | $3.4 \times 10^4$ | 1.3 | $3.3 \times 10^{-4}$ | $1.2 \times 10^{-3}$ |
| (ii) | $5.5 \times 10^{-6}$ | 57.8 | $3.3 \times 10^4$ | 0.47 | $3.5 \times 10^{-5}$ | $1.7 \times 10^{-4}$ |

<sup>(t)</sup> theory and <sup>(m)</sup> measured from experiment or simulation

Given the complexity of the intracellular environment, many factors could influence droplet size and dynamics. It has been proposed that droplet size could be maintained by active processes, such as auto-inhibition of growth (aka “enrichment-inhibition”) [16] or homogeneous chemical conversion of biomolecules between sticky and nonsticky forms [17]. However, our droplet-forming particles are unlikely to be subject to active regulation. Alternatively, experiment and theory have shown that in a synthetic polymer network compressive stresses can frustrate phase separation, control the size of droplets, and reverse the direction of Ostwald ripening [18–20]. These observations raise the question whether the chromatin network could limit droplet growth [11].

To address the role of chromatin in nuclear phase separation, we first perform coarse-grained molecular dynamics (MD) simulations using LAMMPS [21] to investigate droplet formation in a crosslinked chromatin network (Fig. 2). Briefly, the simulation consists of three components (Fig. 2(a)): First, the droplets are composed of particles that attract each other, via a truncated and shifted Lennard-Jones potential:

**FIG. 2.**
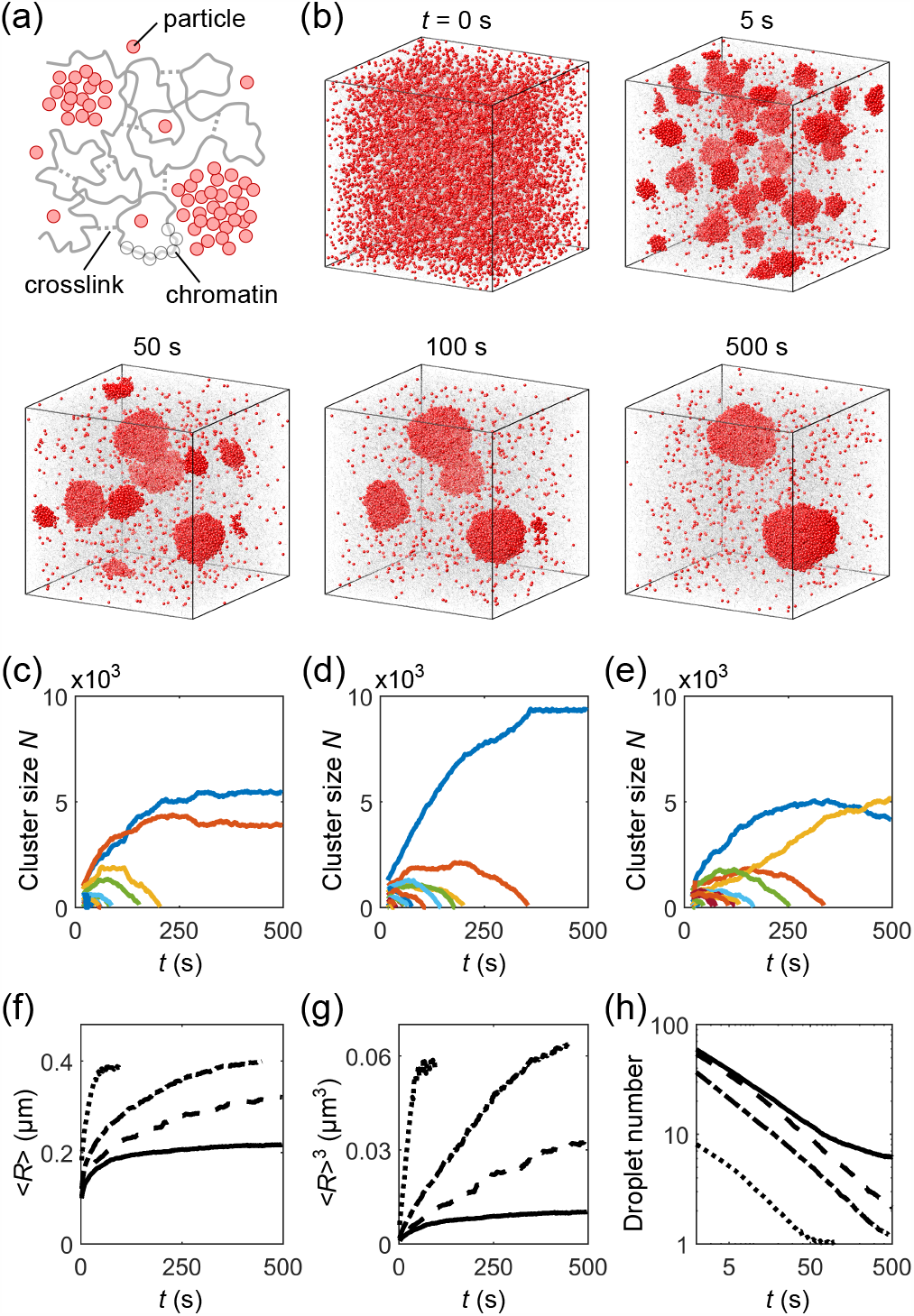
Coarse-grained MD simulation of phase separation of droplet-forming particles in a crosslinked chromatin network. (a) Schematic: particles (red), chromatin (gray, beads and backbone), and crosslinks (gray, dashed lines). (b) Time course of droplets (red particles) embedded in a chromatin network (gray) at crosslink density 7 *µ*M. Chromatin beads are shown at reduced size for better visualization. (c)-(e) Corresponding examples of time evolution of number of particles in individual droplets (colors), where (c) is for the simulation in (b). (f) Mean radius of droplets ⟨*R*⟩, (g) mean radius cubed ⟨*R*⟩^3^, and (h) number of droplets as functions of time *t* for simulations without chromatin (dotted), with chromatin uncrosslinked (dash-dotted), and crosslinked at densities 7 *µ*M (dashed) and 14 *µ*M (solid).

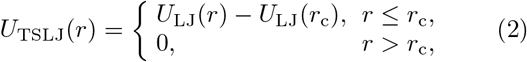

where

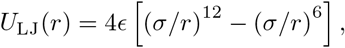

*r* is the distance between particles, *ϵ* = 0.7 *k*_B_*T, σ* = 0.03 *µ*m, and *r*_c_ = 2.5*σ*. The particles alone spontaneously phase separate (Fig. S2) [22]. Second, the chromatin is modeled as a chain of self-avoiding beads connected by soft springs, through a finite extensible nonlinear elastic (FENE) potential [23]:

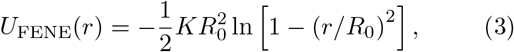

where *R*_0_ = 0.13 *µ*m and 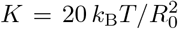. To account for the mechanical elasticity of chromatin, we crosslink the chain randomly, producing a chromatin “gel” network [7,24]. Crosslinks are modeled with the same FENE potential. Third, based on the experimental observation that droplets exclude chromatin [11,12], we introduce a short-ranged repulsion between the phase-separating particles and the chromatin beads. The repulsion is modeled through the LJ potential, Eq. (2), with *ϵ* = 1 *k*_B_*T, σ* = 0.03 *µ*m, and *r*_c_ = 1.12*σ*. Chromatin beads also repel each other via the same LJ potential. We model 10^4^ particles and 10^5^ chromatin beads in a 2 ×2 ×2 *µ*m^3^ simulation box, yielding 18% volume fraction of chromatin. The system evolves according to Langevin dynamics [25] with periodic boundary conditions. For details see [14].

Figure 2(b) shows snapshots of droplets coarsening in a crosslinked chromatin network. The initial configuration consists of particles (red) randomly distributed within the chromatin (gray). To mimic optogenetic activation [11], we turn on attractive interactions between particles at *t* = 0 s. The resulting phase separation involves nucleation of small droplets from the supersaturated bulk solution (*t* = 5 s), followed by droplet ripening and coalescence (*t* = 50, 100 s), and eventually coexistence of multiple droplets (*t* = 500 s). Fig. 2(c)-2(e) shows the number of particles in individual droplets as functions of time for three simulations (with different random crosslinking) at the same overall crosslink density 7 *µ*M (i.e., about 1 crosslink per cube of side length 0.06 *µ*m). (c) is the quantification of the simulation in (b)in which two droplets of different sizes coexist. (d) shows a case where all small droplets evaporate leaving a single large droplet. (e) shows a surprising case in which a larger droplet loses its material to a smaller one, reversing the normal direction of ripening. Such “reverse ripening” has been observed in synthetic polymer systems when a stiff gel containing large droplets is placed next to a soft gel containing small droplets [20]. Clearly, the crosslinked chromatin influences the equilibrium as well as the dynamics of droplet phase separation.

To disentangle the impacts of chromatin *per se* and its crosslinking into a network, we performed a hierarchy of simulations: Case (i) phase separation of droplet-forming particles without chromatin, Case (ii) phase separation in uncrosslinked chromatin, Case (iii) and Case (iv) phase separation in chromatin networks crosslinked at densities 7 *µ*M and 14 *µ*M. Fig. 2(f)-2(h) shows the mean radius of droplets ⟨*R* ⟩, mean radius cubed ⟨*R* ⟩^3^, and number of droplets as functions of time *t* for Cases (i)-(iv), over 50-160 simulations each.

Theoretically, in Cases (i) and (ii) in the absence of merging events, we expect the average radius of droplets ⟨*R* ⟩ to grow according to standard Ostwald ripening [15]:

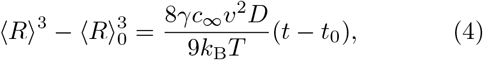

where *t*_0_ is the onset time of ripening, i.e., the end of the initial stage during which droplets grow directly from the supersaturated solution, and ⟨*R* ⟩_0_ = ⟨*R*(*t*_0_) ⟩. The remaining variables have the same definitions as in Eq. (1).

In Case (i) with no chromatin, Eq. (4) predicts *d* ⟨*R* ⟩ ^3^*/dt* = 3.3 ×10^*−*4^ *µ*m^3^*/*s (parameter values in Table I). Fitting the ⟨*R* ⟩ ^3^ versus *t* curve in Fig. 2(g) for Case (i) from *t* = 2 to 20 s yields a higher rate *d* ⟨*R* ⟩^3^*/dt* = 1.2 ×10^*−*3^ *µ*m^3^*/*s. We expect the deviation is due to the droplets occupying a non-negligible volume fraction along with merging events (Fig. S3), both of which are known to speed up droplet growth [12,26]. The rate of increase of ⟨*R* ⟩^3^ slows down at long times due to the finite size of the system, as all simulations end with a single droplet (Fig. 2(h)).

Comparing Cases (i) and (ii), we observe that chromatin acts as a crowder by physically occupying space: this both reduces the dilute-phase concentration threshold *c*_*∞*_ and slows down molecular diffusion *D* (Table I). The decrease in *c*_*∞*_ increases the degree of initial super-saturation and thus reduces the nucleation barrier for droplet formation. As a result, more droplets nucleate from the supersaturated solution (Fig. 2(h)), resulting in smaller droplets at the onset of droplet coarsening. Moreover, the reduction of chromatin polymer entropy near an interface raises the droplet surface free energy [27]. Overall, Eq. (4) predicts *d* ⟨*R* ⟩^3^*/dt* = 3.8 ×10^*−*5^ *µ*m^3^*/*s for Case (ii) and simulation yields *d* ⟨*R* ⟩ ^3^*/dt* = 1.7 ×10^*−*4^ *µ*m^3^*/*s, which is again roughly 4-fold higher, likely due to finite droplet volume fraction and mergers (Fig. S3). Nevertheless, the presence of uncrosslinked chromatin in our simulations slows down droplet growth by ∼10-fold.

In Cases (iii) and (iv) we introduced randomly placed, irreversible crosslinks between chromatin beads separated by long genomic distances [14]. Such long-range crosslinks at a density above the percolation threshold make chromatin gel-like, which can mechanically suppress the growth of large droplets, as these strain the network. The two droplets shown by blue curves in Figs. 2(c) and 2(e) are such examples. In fact, even when the final state is a single droplet (e.g. Fig. 2(d)), we still find a shell of stretched chromatin surrounding the droplet (Fig. 3(a)). Crosslinks also suppress droplet mergers (Fig. S3). Importantly, droplet growth in Case and especially in Case (iv) deviates from the conventional linear dependence on time (Fig. 2(g)). Indeed, the growth of droplets can be completely stalled by the stretched chromatin, leading to the coexistence of multiple droplets – six on average in Case (iv) (Figs. 2(g) and S3).

**FIG. 3.**
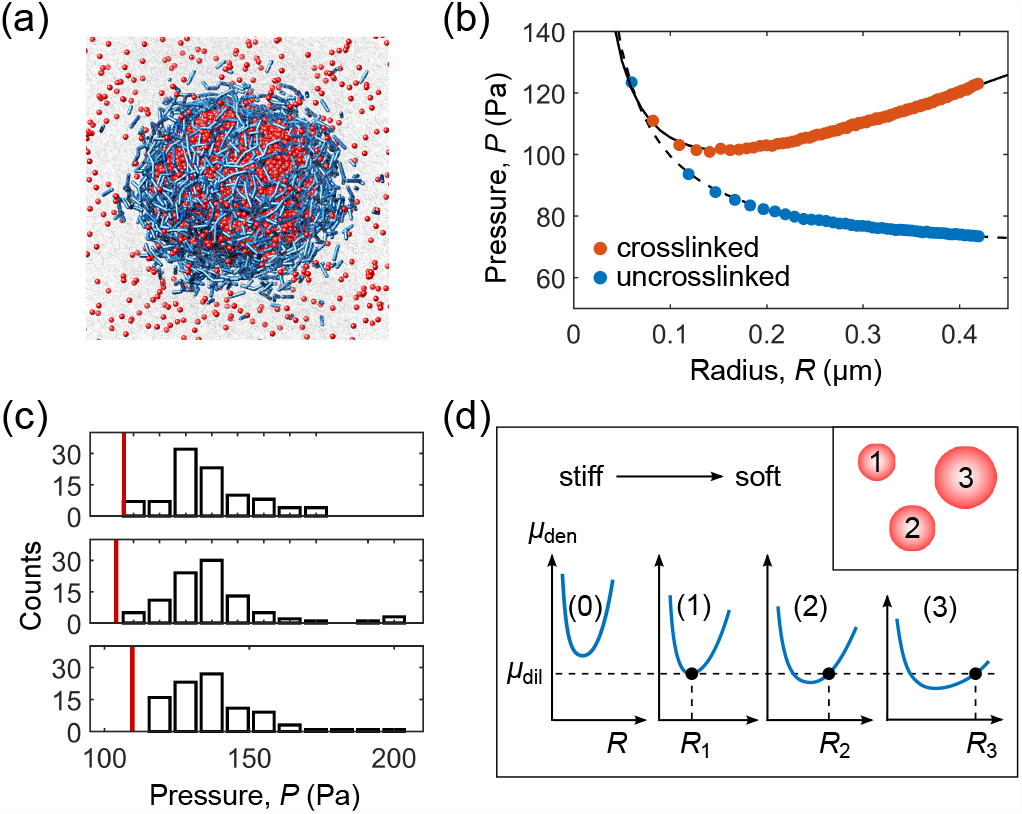
Mechanical interactions with chromatin quantitatively control droplet number, size, and placement. (a) A shell of stretched chromatin backbones and crosslinks (blue) surrounding a droplet (red) in Case (iii). Highlighted bonds are stretched by more than 8% of their mean length prior to droplet formation. (b) Pressure *P* on an inserted sphere of radius *R* in crosslinked chromatin (red) and uncrosslinked chromatin (blue). Solid and dashed curves are fits to Eq. (6). (c)Comparison of pressure by chromatin on a sphere (radius *R* = 0.3 *µ*m) at the actual location of a single large droplet (red vertical line) to that on 95 spheres (histogram) randomly inserted in the same chromatin network, for three examples of Case (iii). (d) Schematic of droplet number, size, and placement control by the variable local stiffness of crosslinked chromatin. Droplet sizes are determined by the position-dependent chemical potential *µ*_den_(*R*_*i*_), which in turn depends on the local crosslink density. No droplet forms at location (0) as stretching the chromatin there is too energetically costly.

To understand how the crosslinked chromatin controls the number, size, and placement of droplets, we consider the equilibrium conditions for droplets (1) temperature balance, (2) pressure balance, and (3) chemical potential balance [28]:

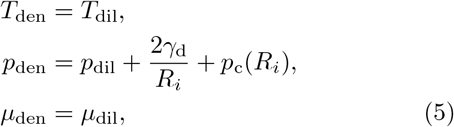

where *γ*_d_ is the intrinsic surface tension of the droplet, *R*_*i*_ is the radius of the *i*^th^ droplet, the term 2*γ*_d_*/R*_*i*_ follows from the Young–Laplace equation [29], and *p*_c_(*R*_*i*_) is the pressure from the chromatin on the *i*^th^ droplet. Because droplets and chromatin repel, there is a clear separation of droplet and chromatin materials at the interface, which allows us to separate their individual contributions to surface energy and hence pressure. Note that the equilibrium equation of state for pure droplet material, *p*_den_ = *p*(*T, µ*), implies that when temperature and chemical potential are balanced, all droplets must have the same internal pressure.

How strongly does chromatin heterogeneity embodied by *p*_c_(*R*_*i*_) influence the final placement of droplets? To systematically measure *p*_c_ as a function of droplet size and location, we place spheres of controllable sizes in different locations. Specifically, in each simulation, we insert two spheres far apart, and vary their sizes while keeping their total volume fixed. (This ensures that the chromatin always takes up the same volume and thus exerts the same pressure.) We record the pressure by chromatin on each sphere as a function of its radius. Fig. 3(b) shows a representative *p*_c_ versus *R* curve in crosslinked chromatin (Case (iii)), compared to uncrosslinked chromatin (Case (ii)). We find that the pressure from uncrosslinked chromatin consists of two parts: a constant bulk pressure *p*_c0_, and a term 2*γ*_c_*/R* from polymer entropy reduction near a curved surface [27]. For crosslinked chromatin, the pressure follows the same trend at small *R* but increases as the sphere grows above *R* ∼ 0.1 *µ*m. This pressure increase identifies the regime where the chromatin begins to elastically constrain droplet growth. We fit the simulation results with

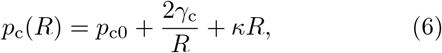

where the linear term *κR* accounts for the strain-stiffening effect of biopolymer networks [30], such as the crosslinked chromatin, which is consistent to the leading order with the hyperelastic Mooney-Rivlin model [31]. For Case (ii) (uncrosslinked chromatin), *κ* = 0 and *p*_c_(*R*) is location independent. By contrast, in crosslinked chromatin, *p*_c_(*R*) depends strongly on location. In Fig. 3(c), we compare the pressure on a sphere at the location where a single large droplet actually formed to randomly located spheres. In all simulations, the sphere at the droplet’s location experienced much lower pressure. We conclude that droplets grow preferentially at places where the mechanical stress from the chromatin is low, consistent with previous experimental results [11,19]. We note that the actual droplets are not always round due to the competition between chromatin stretching and droplet surface tension, but aspherical droplets are also observed in cells [10].

The observed heterogeneity of *p*_c_(*R*_*i*_) leads us to propose a quantitative picture of how crosslinked chromatin controls droplets (Fig. 3(d)). During the initial stage of droplet nucleation, small droplets form in multiple places. As droplets grow large enough to stretch the chromatin, droplets formed in regions with high local crosslink densities have higher internal pressures, whereas droplets formed in less crosslinked regions have lower internal pressures. According to the Gibbs–Duhem equation [32], the chemical potential of the particles in the *i*^th^ droplet is approximately

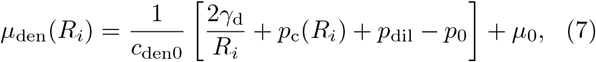

where *c*_den0_, *p*_0_, and *µ*_0_ are the dense-phase concentration, pressure, and chemical potential for phase balance with a flat interface, and we approximate the dense phase to be incompressible. Eq. (7) implies that particles in droplets with higher internal pressures have higher chemical potentials. Thus these droplets will lose particles to droplets with lower internal pressures until every droplet has the same chemical potential as the dilute phase. (As the droplets readjust their sizes, the chemical potential in the dilute phase also gradually decreases.) In Fig. 3(d) we schematically illustrate how droplet sizes are determined by the position-dependent chemical potential *µ*_den_(*R*_*i*_). Finally, the number of droplets that can form depends on the total available material in the system, as the total particle number *N* is conserved:

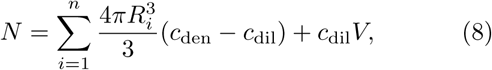

where *n* is the total number of droplets, *c*_den_ and *c*_dil_ are the dense- and dilute-phase concentrations, and *V* the total volume. Motivated by the striking observation that optogenetically-generated droplets in the nucleus can display suppressed coarsening dynamics, we studied the impact of chromatin mechanics on phase separation in the nucleus. We found that the stretching of crosslinked chromatin can mechanically alter droplet evolution as well as quantitatively control the number, size, and placement of droplets. It is thus possible that the observed suppressed coarsening in experiment is due to the stretching of chromatin around the initially generated large droplets.

There remain open questions. How does chromatin’s viscoelasticity (i.e. reversibility of crosslinks) impact phase separation? In which parameter regions does ripening verus merging dominate coarsening dynamics [12], and where do specific biomolecular condensates (e.g. nucleoli, PML bodies) sit in this parameter space? Interestingly, many biomolecules, such as transcription factors [33], heterochromatin protein 1 [34], and BRD4 [35], have an affinity for chromatin. How does phase separation proceed if the interaction between droplets and chromatin is attractive instead of repulsive? We hope the results presented here will encourage future work on such questions.

## ACKNOWLEDGMENTS

We thank Edgar Blokhuis and Ben Weiner for insightful suggestions. This work was supported in part by the National Science Foundation, through the Center for the Physics of Biological Function (PHY-1734030), the Graduate Research Fellowship Program (DCE-1656466, D.S.W.L.), NIH Grants R01 GM082938, U01 DA040601, and the Howard Hughes Medical Institute.

## Notes

### Competing Interest Statement

The authors have declared no competing interest.

## References

[1] Y. Shin and C. P. Brangwynne, Science 357, eaaf4382 (2017).

[2] S. F. Banani, H. O. Lee, A. A. Hyman, and M. K. Rosen, Nat. Rev. Mol. Cell Biol. 18, 285 (2017).

[3] C. P. Brangwynne, C. R. Eckmann, D. S. Courson, A. Rybarska, C. Hoege, J. Gharakhani, F. Jülicher, and A. A. Hyman, Science 324, 1729 (2009).

[4] A. Molliex, J. Temirov, J. Lee, M. Coughlin, A. P. Kana-garaj, H. J. Kim, T. Mittag, and J. P. Taylor, Cell 163, 123 (2015).

[5] E. S. F. Rosenzweig, B. Xu, L. K. Cuellar, A. Martinez-Sanchez, M. Schaffer, M. Strauss, H. N. Cartwright, P. Ronceray, J. M. Plitzko, F. Forster, et al., Cell 171, 148 (2017).

[6] C. P. Brangwynne, P. Tompa, and R. V. Pappu, Nat. Phys. 11, 899 (2015).

[7] A. D. Stephens, E. J. Banigan, S. A. Adam, R. D. Gold-man, and J. F. Marko, Mol. Biol. Cell 28, 1984 (2017).

[8] A. D. Stephens, E. J. Banigan, and J. F. Marko, Curr. Opin. Cell Biol. 58, 76 (2019).

[9] V. Pareek, H. Tian, N. Winograd, and S. J. Benkovic, Science 368, 283 (2020).

[10] M. Feric, N. Vaidya, T. S. Harmon, D. M. Mitrea, L. Zhu, T. M. Richardson, R. W. Kriwacki, R. V. Pappu, and C. P. Brangwynne, Cell 165, 1686 (2016).

[11] Y. Shin, Y.-C. Chang, D. S. Lee, J. Berry, D. W. Sanders, P. Ronceray, N. S. Wingreen, M. Haataja, and C. P. Brangwynne, Cell 175, 1481 (2018).

[12] D. S. Lee, N. S. Wingreen, and C. P. Brangwynne, bioRxiv (2020).

[13] D. Bracha, M. T. Walls, M.-T. Wei, L. Zhu, M. Kurian, J. L. Avalos, J. E. Toettcher, and C. P. Brangwynne, Cell 175, 1467 (2018).

[14] See Supplemental Material at … for additional information about experiment, simulation, and theory.

[15] I. M. Lifshitz and V. V. Slyozov, J. Phys. Chem. Solids 19, 35 (1961).

[16] J. Söding, D. Zwicker, S. Sohrabi-Jahromi, M. Boehning, and J. Kirschbaum, Trends Cell Biol. 30, 4 (2020).

[17] J. D. Wurtz and C. F. Lee, Phys. Rev. Lett. 120, 078102 (2018).

[18] R. W. Style, T. Sai, N. Fanelli, M. Ijavi, K. Smith-Mannschott, Q. Xu, L. A. Wilen, and E. R. Dufresne, Phys. Rev. X 8, 011028 (2018).

[19] K. A. Rosowski, T. Sai, E. Vidal-Henriquez, D. Zwicker, R. W. Style, and E. R. Dufresne, Nat. Phys. 16, 422 (2020).

[20] K. Rosowski, E. Vidal-Henriquez, D. Zwicker, R. Style, and E. R. Dufresne, Soft Matter (2020).

[21] S. Plimpton, J. Comput. Phys. 117, 1 (1995).

[22] A. E. van Giessen and E. M. Blokhuis, J. Chem. Phys. 131, 164705 (2009).

[23] K. Kremer and G. S. Grest, J. Chem. Phys. 92, 5057 (1990).

[24] N. Khanna, Y. Zhang, J. S. Lucas, O. K. Dudko, and C. Murre, Nat. Commun. 10, 1 (2019).

[25] P. Langevin, C. R. Acad. Sci. 146, 530 (1908).

[26] J. H. Yao, K. Elder, H. Guo, and M. Grant, Physica A 204, 770 (1994).

[27] P. G. de Gennes, C. R. Acad. Sci. Paris B 288, 359 (1979).

[28] L. D. Landau and E. M. Lifshitz, Statistical Physics (Course of Theoretical Physics, Volume 5) (Elsevier, 2013).

[29] J. S. Rowlinson and B. Widom, Molecular theory of capillarity (Courier Corporation, 2013).

[30] K. A. Erk, K. J. Henderson, and K. R. Shull, Biomacromolecules 11, 1358 (2010).

[31] M. Kothari and T. Cohen, arXiv preprint 2004.13238 (2020).

[32] P. Perrot, A to Z of Thermodynamics (Oxford University Press, 1998).

[33] A. Boija, I. A. Klein, B. R. Sabari, A. Dall’Agnese, E. L. Coffey, A. V. Zamudio, C. H. Li, K. Shrinivas, J. C. Manteiga, N. M. Hannett, et al., Cell 175, 1842 (2018).

[34] A. G. Larson, D. Elnatan, M. M. Keenen, M. J. Trnka, J. B. Johnston, A. L. Burlingame, D. A. Agard, S. Redding, and G. J. Narlikar, Nature 547, 236 (2017).

[35] B. A. Gibson, L. K. Doolittle, M. W. Schneider, L. E. Jensen, N. Gamarra, L. Henry, D. W. Gerlich, S. Redding, and M. K. Rosen, Cell 179, 470 (2019).

