## Supplemental Material for "Mechanical frustration of phase separation in the cell nucleus by chromatin"

### Supplementary Material for Zhang *et al.*

#### I. EXPERIMENTAL METHODS

We apply the Corelet system [1] to generate synthetic liquid nuclear bodies in living cells. Briefly, the Corelet system is comprised of two components: a ferritin "core", consisting of ferritin subunits, which oligomerize into a 24-mer, fused to a green fluorescent protein (eGFP) and an iLiD, plus an intrinsically disodered region (IDR) fused to mCherry (mCh) fluorescent protein and an sspB domain. Upon "activation" of the sspB domain by exposure to blue light, the iLiD and sspB heterodimerize, resulting in the decoration of core molecules with multiple IDRs which effectively oligomerizes the IDRs, triggering phase separation.

This optogenetic approach allows for spatiotemporal control of phase separation inside cells using patterned blue light stimulation. By focusing the light on specific parts of the nucleus, we can force these particular parts of the nucleus deep into the two-phase region of the phase diagram [1]. We can thus generate large droplets using patterned local activation, before switching to global activation, which drives the phase separation in the entire nucleus, Fig. S1(a)-(b). Due to a diffusive capture mechanism [1], the large droplets generated by local activation initially have higher IDR-to-core ratios than the small droplets generated by global activation. As a result, the areas of the large droplets decrease transiently while this ratio equilibrates, as shown in Fig. S1(c). The time courses of droplet sizes in Fig. S1(c) suggest that equilibration of the IDR-to-core ratio is completed around 60 min; we therefore focus in the main text on the time evolution of droplet sizes starting at 60 min after global activation.

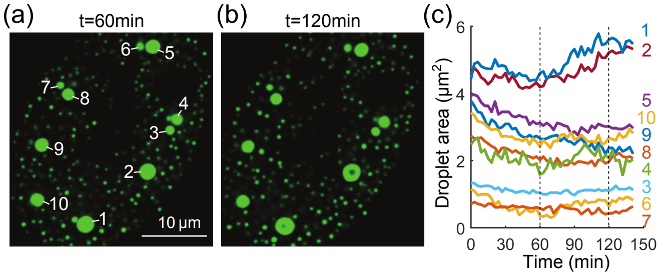


FIG. S1. Time evolution of optogenetically-generated fluorescent protein droplets in the nucleus of a U2OS cell. (a) Snapshot of droplets at 60 min after global activation. (b) Snapshot at 120 min. (c) Time evolution of areas of large droplets labeled in (a).

##### A. Cell culture

U2OS (a kind gift from Mark Groudine lab, Fred Hutchinson Cancer Research Center) and Lenti-X 293T (Takara) cells were cultured in growth medium consisting of Dulbecco's modified Eagle's medium (GIBCO), 10 percent fetal bovine serum (Atlanta Biologicals), and 10 U/mL Penicillin-Streptomycin (GIBCO), and incubated at 37 °C and 5% $CO_2$ in a humidified incubator.

##### B. Lentiviral transduction

Lentivirus was produced by transfecting the transfer plasmid, pCMV-dR8.91, and pMD2.G (9:8:1, mass ratio) into Lenti-X cells grown to approximately 80 percent confluence in 6-well plates using FuGENE HD Transfection Reagent (Promega) per manufacturer's protocol. A total of 3 $\mu$g plasmid and 9 $\mu$L of transfection reagent were delivered into each well. After 60-72 hours, supernatant containing viral particles was harvested and filtered with 0.45 $\mu$m filter (Pall Life Sciences). Supernatant was immediately used for transduction or aliquoted and stored at $-80$ °C. U2OS were seeded at 10 percent confluence in 96-well plates and 20-200 $\mu$L of filtered viral supernatant was added to the cells. Media containing virus was replaced with fresh growth medium 24 hr post-infection. Infected cells were imaged no earlier than 72 hr after infection.

##### C. Microscopy

All images were taken with a spinning disk (Yokogawa CSU-X1) confocal microscope with a 100X oil immersion Apo TIRF objective (NA 1.49) and an Andor DU-897 EMCCD camera on a Nikon Eclipse Ti body, with a 488nm laser for imaging GFP, and a 561nm laser for imaging mCherry. The imaging chamber was maintained at 37 °C and 5% $CO_2$ (Okolab) with a 96-well plate adaptor. Both local and global activation were performed by constantly stimulating the sample with blue light (485nm) driven by a Lumencor SpectraX light engine and patterned using a Mightex Polygon digital micromirror device (DMD) controlled by Nikon Elements software. Local activation was performed for 10 minutes, and global activation was subsequently performed for 2.5 hrs, during which images were taken every 3 minutes in single central $z$-plane for each nucleus, using Perfect Focus to maintain focal depth of the sample over time.

##### D. Estimation of ripening rate

We consider a large droplet of radius $R_1$ surrounded by many small droplets of size $R_2$ all at a distance $L$. The concentration field $c(\vec{r},t)$ outside the droplets satisfies the diffusion equation:

$$\frac{\partial c(\vec{r},t)}{\partial t} = D\nabla^2 c(\vec{r},t), \tag{S1}$$

where $D$ is the diffusion coefficient of the molecule in the dilute phase.

We assume the concentrations at the boundaries of the droplets are set by the Gibbs-Thomson relation [2]:

$$\begin{aligned} c(r=R_1,t) &= c_\infty\left(1+\frac{2\gamma v}{k_\mathrm{B}TR_1}\right), \\ c(r=L,t) &= c_\infty\left(1+\frac{2\gamma v}{k_\mathrm{B}TR_2}\right), \end{aligned} \tag{S2}$$

where $c_\infty$ is the dilute concentration threshold, $\gamma$ is the surface tension, and $v$ is the molecular volume in the dense phase. Note that the second line arises simply because by assumption there are many droplets of size $R_2$ at $r=L$.

To determine the flux into the large droplet, we first find the quasi-steady state solution to Eq. (S1) (i.e. $\nabla^2 c_\mathrm{ss}(\vec{r})=0$) with respect to the boundary conditions in Eq. (S2),

$$c_\mathrm{ss}(r) = c_\infty\left[1+\frac{2\gamma v}{k_\mathrm{B}T}\left(\frac{1}{L}\frac{\frac{L}{R_2}-1}{1-\frac{R_1}{L}} - \frac{1}{r}\frac{\frac{R_1}{R_2}-1}{1-\frac{R_1}{L}}\right)\right]. \tag{S3}$$

The flux across the boundary of the central droplet is:

$$J = 4\pi R_1^2 D\left.\frac{dc_\mathrm{ss}}{dr}\right|_{r=R_1} = \frac{8\pi\gamma c_\infty v D}{k_\mathrm{B}T}\frac{\frac{R_1}{R_2}-1}{1-\frac{R_1}{L}}. \tag{S4}$$

Finally, as the change in the volume of the droplet over time is given by the flux of material into the droplet, $dV_1/dt = vJ$, we obtain Eq. (1) in the main text:

$$\frac{dR_1^3}{dt} = \frac{6a\gamma c_\infty v^2 D}{k_B T}, \tag{S5}$$

where $a=(R_1/R_2-1)/(1-R_1/L)$ is a geometrical factor.

The parameters used to estimate the theoretically predicted ripening rate for the experimental conditions are: $v = 6.7\times10^4\,\mathrm{nm}^3$, $c_\infty = 2\,\mu\mathrm{M}$, $D_\mathrm{core} = 3\,\mu\mathrm{m}^2/\mathrm{s}$, and $D_\mathrm{IDR} = 44\,\mu\mathrm{m}^2/\mathrm{s}$ as measured in [1]. We take $D\approx 1\,\mu\mathrm{m}^2/\mathrm{s}$ for the bound complex of core and IDR and take $\gamma\approx 5\times10^{-7}\,\mathrm{N/m}$ from [3].

#### II. SIMULATION METHODS

We performed coarse-grained molecular dynamics (MD) simulations using LAMMPS [4] to investigate the impact of chromatin mechanics on phase separation in the cell nucleus. A hierarchy of simulations were designed to disentangle the roles of chromatin and its crosslinking into a network: Case (i) phase separation of droplet-forming particles without chromatin, Case (ii) phase separation in uncrosslinked chromatin, Case (iii) and Case (iv) phase separation in chromatin networks crosslinked at densities $7\,\mu$M and $14\,\mu$M, respectively.

##### A. Case (i)

The simulation consists of particles that interact with each other via a truncated and shifted Lennard-Jones potential:

$$U_\mathrm{TSLJ}(r) = \begin{cases} U_\mathrm{LJ}(r) - U_\mathrm{LJ}(r_\mathrm{c}), & r\le r_\mathrm{c}, \\ 0, & r > r_\mathrm{c}, \end{cases} \tag{S6}$$

where

$$U_\mathrm{LJ}(r) = 4\epsilon\left[\left(\frac{\sigma}{r}\right)^{12} - \left(\frac{\sigma}{r}\right)^6\right],$$

and $r$ is the distance between particles. 160 simulation replicates were performed, each contained $10^4$ pariticles in a box of size $2\,\mu\mathrm{m}\times2\,\mu\mathrm{m}\times2\,\mu\mathrm{m}$ with periodic boundary conditions. The system evolves according to Langevin dynamics [5] in the $NVT$ ensemble at $T=300\,\mathrm{K}$:

$$m\frac{d^2\vec{r}_i}{dt^2} = -\gamma\frac{d\vec{r}_i}{dt} - \nabla_{\vec{r}_i}U(\vec{r}_1,...,\vec{r}_N) + \vec{f}, \tag{S7}$$

where $\vec{r}_i$ is the coordinate of particle $i$, $m$ is its mass, $\gamma$ is the friction coefficient, $\vec{f}$ is random thermal noise, and the potential $U(\vec{r}_1,...,\vec{r}_N)$ contains all interactions between particle $i$ and other particles. This Langevin thermostat is applied using a damping factor $\tau = m/\gamma = 250\,\mu$s, mass of particle $m = 706.9\,\mathrm{pg}$, and integration step size $dt = 5\,\mu$s. These parameters give the particle the right diffusion coefficient $D = k_\mathrm{B}T/(3\pi\eta d)$ at times much longer than $\tau$, where $\eta = 0.01\,\mathrm{kg/m/s}$ is the viscosity of nucleoplasm (assumed ∼10 times that of water) and $d = 0.03\,\mu$m the diameter of the particle. For initial equilibration, the interactions between particles are set to be purely repulsive with $\epsilon = 1\,k_\mathrm{B}T$, $\sigma = 0.03\,\mu$m, and $r_\mathrm{c} = 2^{1/6}\sigma$ in Eq. (S6). The system is equilibrated for $10^7$ steps. To mimic optogenetic activation, attractive interactions are turned on at $t = 0\,$s by changing the interaction parameters in Eq. (S6) to $\epsilon = 0.7\,k_\mathrm{B}T$, $\sigma = 0.03\,\mu$m, and $r_\mathrm{c} = 2.5\sigma$. The positions of all particles are recorded every $4\times10^5$ steps for 50 recordings in each simulation replicate. Clusters (droplets) are identified using Matlab function *knnsearch* with a cutoff distance $0.05\,\mu$m, i.e., any two particles with a center-to-center distance less than $0.05\,\mu$m are considered to be part of the same cluster. All simulations end with a single dense droplet surrounded by a dilute solution (Fig. S2). The dilute- and dense-phase concentrations are $0.58\,\mu$M and $48.7\,\mu$M (or 0.0094 and 0.792 in the reduced unit $c^* = c\sigma^3$) consistent with previously reported values 0.0091 and 0.796 [6].

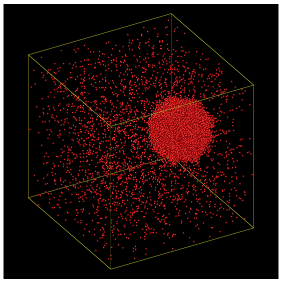

FIG. S2. A single dense droplet formed surrounded by a dilute particle solution for all simulations in Case (i).

#### B. Case (ii)

Here, we add uncrosslinked chromatin chains to the simulation in Case (i). The chromatin is modeled as a chain of self-avoiding beads connected by soft springs, through a finite extensible nonlinear elastic (FENE) potential [7]:

$$U_{\mathrm{FENE}}(r) = -\frac{1}{2}KR_0^2 \ln\left[1-\left(\frac{r}{R_0}\right)^2\right], \qquad \text{(S8)}$$

where $R_0 = 0.13\,\mu$m and $K = 20\,k_\mathrm{B}T/R_0^2$. 110 simulation replicates were performed, each contained $10^4$ particles and $10^5$ chromatin beads (8 chains each with 12500 beads) in a box of size $2\,\mu\mathrm{m} \times 2\,\mu\mathrm{m} \times 2\,\mu\mathrm{m}$ with periodic boundary conditions. The system is equilibrated for $10^7$ steps, using a Langevin thermostat in the $NVT$ ensemble at $T = 300$K with purely replusive interactions for particle-particle, particle-chromatin, and chromatin-chromatin pairs ($\epsilon = 1\,k_\mathrm{B}T$, $\sigma = 0.03\,\mu$m, and $r_\mathrm{c} = 2^{1/6}\sigma$ in Eq. (S6)). Attractions between particles are turned on at time $t = 0$ s by changing the interaction parameters for the particle-particle pairs to $\epsilon = 0.7\,k_\mathrm{B}T$, $\sigma = 0.03\,\mu$m, and $r_\mathrm{c} = 2.5\sigma$. Positions of all particles are recorded every $4 \times 10^5$ steps for 500 recordings.

#### C. Case (iii) and Case (iv)

To account for the mechanical elasticity of chromatin, we further crosslink the chromatin to produce a "gel" network. Briefly, a snapshot of uncrosslinked chromatin chains from Case (ii) is recorded. Linkable bead pairs are identified if (1) the two beads are separated by more than 5 beads away along the chain and (2) the distance between the two beads are within $0.08\,\mu$m. We selected $3.5 \times 10^4$ and $7 \times 10^4$ pairs (i.e., at densities $7\,\mu$M and $14\,\mu$M) from all linkable pairs for Case (iii) and Case (iv). The selection procedure ensures that two selected pairs with bead indices $(i, j)$ and $(i', j')$ are well separated along the chain, i.e., $\sqrt{|i-i'|^2 + |j-j'|^2} > 5$, and is otherwise random. Crosslinks are modeled with the same FENE potential, Eq. (S8). The equilibration and data recording procedures are the same as for Case (ii), except 60 and 50 simulation replicates for Case (iii) and Case (iv) were performed.

Figure S3 shows representative examples of time evolution of number of particles in individual droplets for Cases (ii) to (iv). Comparing Cases (i) and (ii) to Cases (iii) and (iv) reveals that crosslinks suppress droplet coarsening.


[1] D. Bracha, M. T. Walls, M.-T. Wei, L. Zhu, M. Kurian, J. L. Avalos, J. E. Toettcher, and C. P. Brangwynne, Cell **175**, 1467 (2018).
[2] W. Thomson, Proc. R. Soc. Edinb. **7**, 63 (1872).
[3] M. Feric, N. Vaidya, T. S. Harmon, D. M. Mitrea, L. Zhu, T. M. Richardson, R. W. Kriwacki, R. V. Pappu, and C. P. Brangwynne, Cell **165**, 1686 (2016).
[4] S. Plimpton, J. Comput. Phys. **117**, 1 (1995).
[5] P. Langevin, C. R. Acad. Sci. **146**, 530 (1908).
[6] J. Vrabec, G. K. Kedia, G. Fuchs, and H. Hasse, Mol. Phys. **104**, 1509 (2006).
[7] K. Kremer and G. S. Grest, J. Chem. Phys. **92**, 5057 (1990).

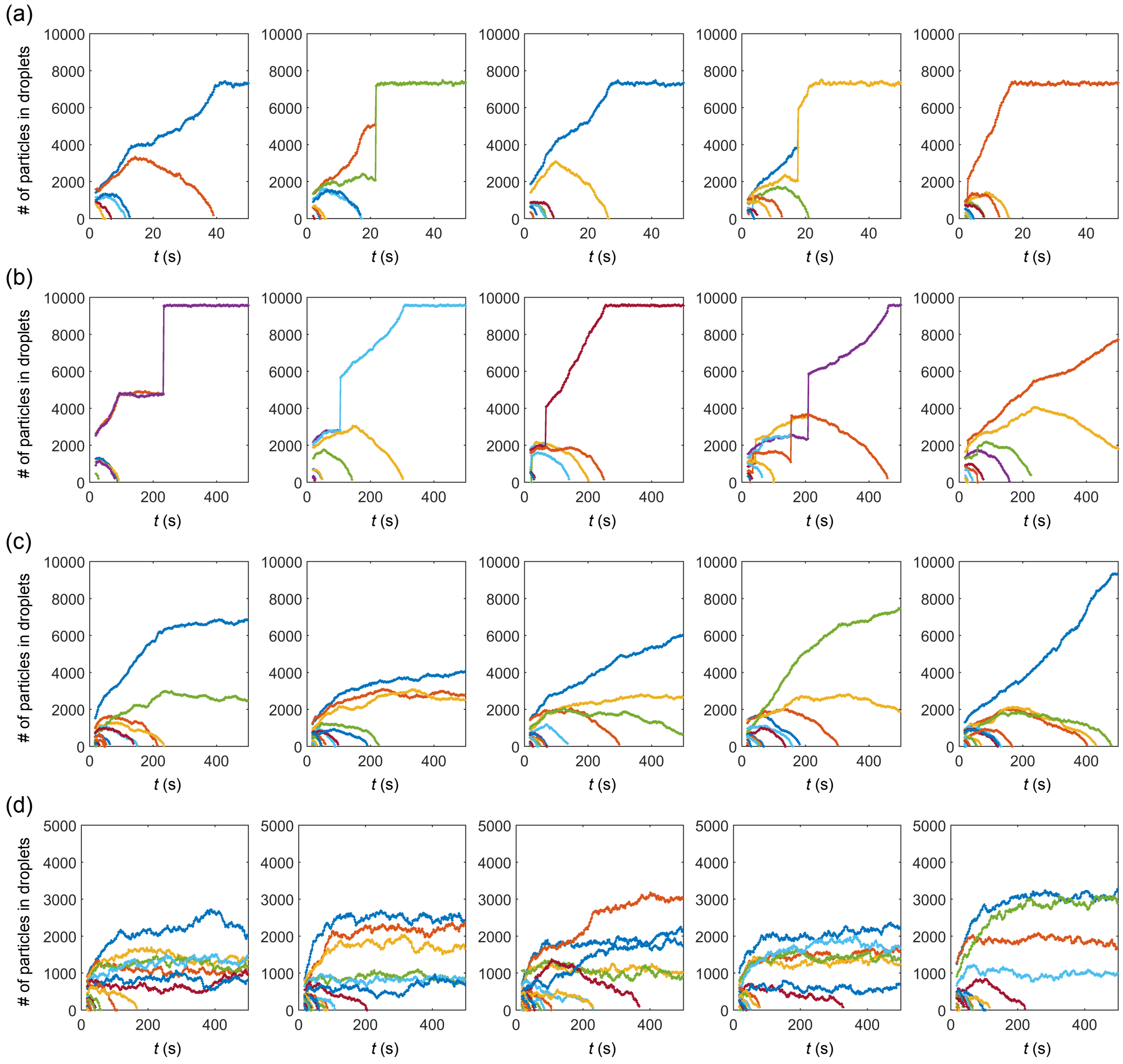


FIG. S3. Crosslinked chromatin suppresses droplet ripening and merging. (a)-(d) Time evolution of number of particles in individual droplets for Cases (i)-(iv).
